# Temporal tracking of Synaptobrevin-1 trafficking reveals SAM-4/BORC-dependent trafficking routes in *C. elegans* neurons

**DOI:** 10.64898/2026.04.29.721573

**Authors:** Badal Singh Chauhan, Aryaman Kunwar, Sandhya P. Koushika

## Abstract

Synaptic vesicle proteins (SVPs) are synthesised in the neuronal soma trafficked as precursor synaptic vesicles (pre-SVs) on route to synapses. While pre-SVs are known to have heterogeneous protein composition and can co-traffic with lysosomal proteins. In this study, we assess the trafficking routes and kinetics of Synatobrevin-1 (SNB-1) released from the ER using the RUSH system *in vivo* in *C. elegans* touch receptor neurons. We showed that ER-released SNB-1 follows at least two temporally distinct trafficking routes. A predominantly anterogradely moving population of SNB-1 carrying vesicles appeared early, within 20 minutes of ER release in the axon without overlap with lysosomal proteins. Another SNB-1 population at 45 minutes post-ER release overlapped with endolysosomal compartments in both the cell body and the axon. Early SNB-1 carrying vesicles co-migrate with a transmembrane synaptic vesicle protein Synaptogyrin (SNG-1) and RAB-27 but fewer with RAB-3, suggesting that SVPs can be co-sorted into the same carriers prior to overlap with lysosomal proteins. The SV–lysosomal protein overlap occurs even when SNB-1 endocytosis on the plasma membrane is reduced in *unc-11/ap180* mutants. Finally, we identified SAM-4/Myrlysin, a subunit of the BORC complex, as a regulator of both the trafficking kinetics of Synaptobrevin-1 intermediates and the cargo composition of pre-SVs. Loss of SAM-4 accelerated SV–lysosomal protein overlap and reduced co-transport of SNG-1 with SNB-1 in early pre-SVs in the axon. Together, these findings reveal heterogeneity in pre-SV biogenesis routes and identify SAM-4 as a key regulator of both the kinetics and cargo composition of synaptic vesicle precursors.

## Introduction

Neurons communicate by releasing neurotransmitters from presynaptic terminals (1). These neurotransmitters are stored in synaptic vesicles (SVs), which contain a characteristic set of membrane proteins known as synaptic vesicle proteins (SVPs) (2, 3). Many SV membrane proteins, including transmembrane proteins and peripheral membrane proteins, are assembled into transport carriers in the neuronal soma; these organelles are called precursors of synaptic vesicles (pre-SVs) and are transported along axons to presynaptic sites (4–7).

Literature from neuroendocrine cells and hippocampal neurons suggests that SV proteins (e.g., synaptophysin and synaptotagmin), after sorting from the trans-Golgi network (TGN) compartment, may transit to the plasma membrane and undergo internalisation (8–12). However, it remains unclear whether this route accounts for the entire pool of newly synthesised SV proteins or only a fraction of them.

Multiple lines of evidence indicate that pre-SVs are compositionally heterogeneous: only subsets of SVPs are transported in the same transport carrier, suggesting multiple pre-SV pools of vesicles present in the axon (5, 6, 13). Notably, SVPs also co-transport with lysosomal proteins in both the soma and axons across multiple model systems (14–18). This SV-lysosomal protein overlap could arise from either biosynthetic trafficking from the cell body or degradative pathways involving used SV proteins. A recent study in hippocampal neurons suggests that SV-endolysosomal intermediate compartments could arise from the biosynthetic routes (19).

Genetic studies have identified several early steps of pre-SV biogenesis; for instance, UNC-16/JIP3 regulates the exclusion of Golgi proteins from pre-SVs (6). Acting within the same genetic pathway, LRK-1/LRRK2 regulates vesicle composition and size via the adaptor protein complexes AP-3 and AP-1, respectively (6). Lysosomes and synaptic vesicles share several regulators involved in the biogenesis and trafficking of these compartments; notably, RAB-2, acting at the Golgi, regulates the export and sorting of SV and lysosomal proteins (20), and loss of LRK-1/LRRK2, APB-3/AP-3, or SAM-4/Myrlysin results in increased overlap between SV and lysosomal proteins (14, 17, 21–24).

The BORC (BLOC-1-Related complex) is a key regulator of the distribution of SV (25) and lysosomal proteins (26). In mammalian neurons, BORC is required for the axonal transport of lysosomal proteins, but not for synaptic vesicle trafficking (18, 26). In contrast, studies in *C. elegans* neurons indicate that BORC is required for the distribution and segregation of both synaptic vesicles and lysosomal proteins (21, 25, 27). Together, these observations suggest that the role of BORC in axonal transport is context-dependent, with distinct effects on synaptic vesicle precursor (SVP) trafficking and lysosomal positioning across different model systems.

In addition to adaptor complexes and BORC, peripheral membrane proteins, such as RAB GTPases, regulate distinct aspects of vesicle trafficking and fusion (28, 29). RAB-3 and RAB-27 are associated with SVs and dense core vesicles (DCVs) and play a role in docking and exocytosis of these vesicles at synapses (30–34). Understanding when these RAB proteins are recruited to newly formed vesicles can provide insights into the maturation process of SVs and their relationship to other secretory organelles.

The compositional heterogeneity of pre-SVs, the co-trafficking of SVPs with lysosomal proteins, and the sharing of common regulators collectively suggest that multiple pathways may deliver SVPs to the axon. Distinguishing between these potential routes is challenging using steady-state imaging, which cannot temporally resolve newly synthesised cargo from the pre-existing pools. Similarly, whether SAM-4/BORC acts on the kinetics of SV–lysosomal intermediate formation, the protein composition of pre-SVs, or both, cannot be resolved without temporally precise tracking of newly synthesised SVPs. The RUSH (Retention Using Selective Hooks) system has emerged as a powerful tool for investigating the kinetics of protein trafficking with temporal precision (35, 36). In hippocampal neurons and DA9 neurons of *C. elegans*, this system reveals that newly synthesised SV proteins are selectively transported to the axon (36).

The precise trafficking routes and kinetics of SVPs during pre-SV biogenesis in vivo remain unexplored. To address this, we combined the RUSH system with live imaging in *C. elegans* mechanosensory PLM neurons to dissect the trafficking routes of newly synthesised SNB-1 from the ER to the axon. Here, we show that ER-released SNB-1 can follow at least two distinct trafficking routes: a faster route that does not go through compartments containing lysosomal proteins and a slower route where SNB-1 overlaps with lysosomal proteins. We further show that RAB-3 and RAB-27 are recruited to ER-released SNB-1-containing vesicles with distinct kinetics. Finally, we identify SAM-4 as a regulator of both the timing of SV–lysosomal intermediate formation and the protein composition of pre-SVs entering the axon, likely acting at the Golgi. Together, these findings reveal heterogeneity in SV biogenesis routes and identify SAM-4 as a key regulator of both the kinetics and protein composition of pre-SVs, advancing our understanding of pre-SV biogenesis in vivo.

## Results

### SNB-1 containing vesicles coming from the biosynthetic route co-migrate with SV and lysosomal proteins at different time points

Synaptic vesicle proteins, including SNB-1/Synaptobrevin and SNG-1/Synaptogyrin, co-migrate with lysosomal proteins in both the cell body and axon (15–17, 21), but the trafficking routes and kinetics taken by SVPs during pre-SV biogenesis remain unclear.

To investigate the trafficking routes of SNB-1, a synaptic vesicle V-SNARE protein in vivo, we use the RUSH system (Retention Using Selective Hooks) to retain SNB-1 in the endoplasmic reticulum (ER). Here, SNB-1 is tagged with a streptavidin-binding peptide (SBP) and the fluorescent protein eGFP, together forming the reporter complex. Streptavidin is tagged with the ER retention signal KDEL, acting as a hook to selectively retain SNB-1::SBP::eGFP in the ER (Fig 1A). This fusion protein is expressed in the touch receptor neurons (TRNs) of *C. elegans*. This permits the synchronised release of SNB-1 upon feeding the organism with biotin, which displaces SNB-1::SBP::eGFP from Streptavidin. This construct can provide SNB-1 function, as shown by the rescue of locomotion defects in a *snb-1* partial loss-of-function mutant (S1B). As the SV proteins require the kinesin-3 motor KIF1A/UNC-104 to exit the cell body (25, 37), synaptic localization of SBP::eGFP tagged SNB-1 is likewise dependent on UNC-104/KIF1A (S1D).

**Figure 1:**
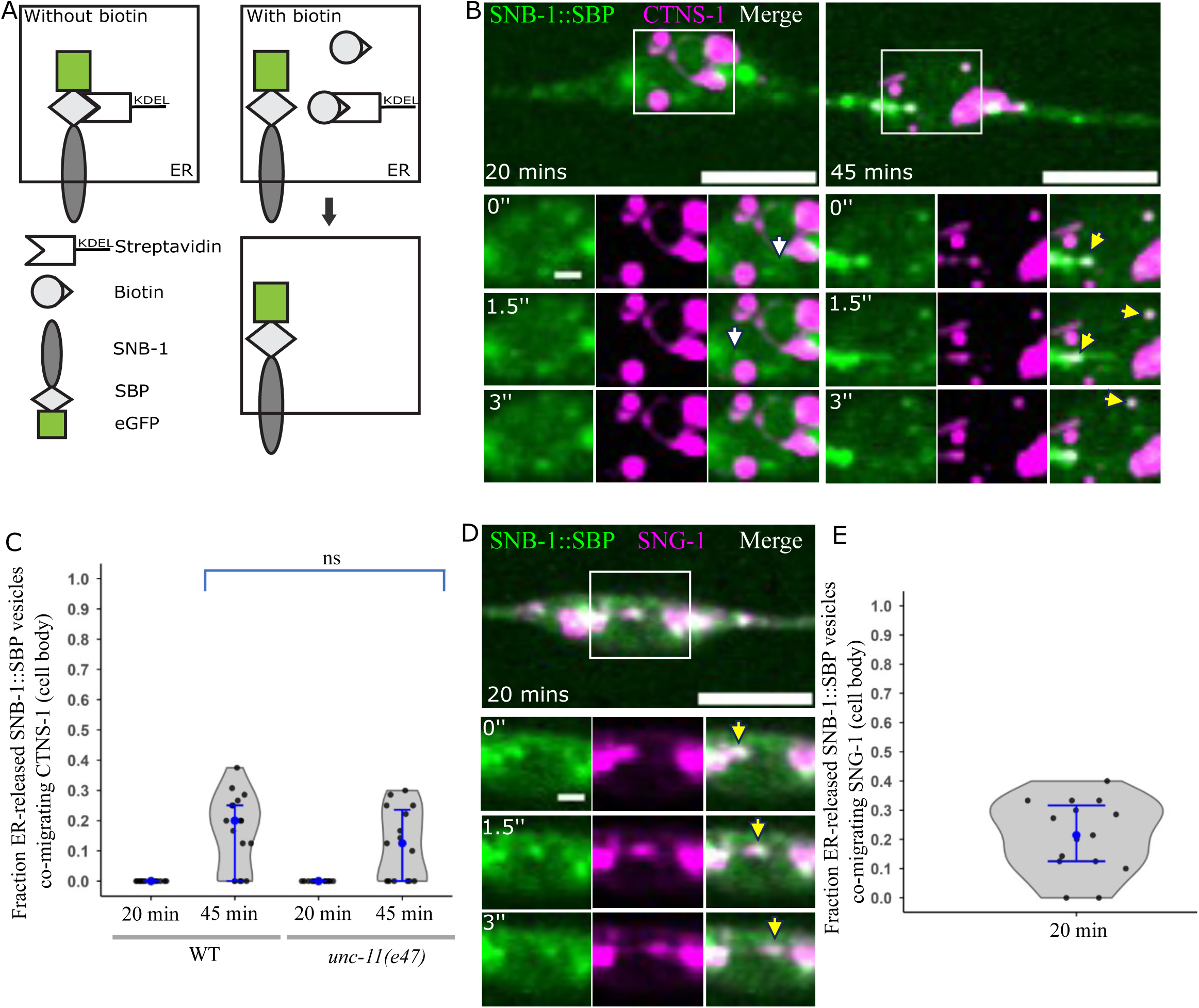
ER-released SNB-1 exhibits temporal separation from lysosomal proteins in the soma. A) Schematics of the RUSH system-before and after biotin feeding, along with its two components Hook (Streptavidin-KDEL) and reporter (SNB-1::SBP::eGFP). B) Representative image of ER-released SNB-1::SBP::eGFP (Green) and CTNS-1::mCherry (Magenta) dynamic compartments after 20 min (left panel) and 45 min (right panel) post-biotin feeding in the PLM cell body. The white arrow indicates ER-released SNB-1::SBP::eGFP containing dynamic compartments lacking CTNS-1::mCherry. The yellow arrow indicates ER-released SNB-1::SBP::eGFP containing dynamic compartments containing CTNS-1::mCherry. Scale bar 5 µm (large panel with ROI), 1 µm (time series panels). C) Violin plot showing the fraction of ER-released SNB-1::SBP::eGFP containing vesicles co-migrating with CTNS-1::mCherry in the cell body of PLM neuron at 20 min and 45 min post-biotin feeding in WT and *unc-11(e47)* mutant animals. Number of animals ≥ 15, Number of SNB-1::SBP::eGFP compartments ≥ 82. WT 45 min post biotin feeding Vs *unc-11(e47)* 45 min post biotin feeding, Mann-Whitney Test, *p*-value= 5.24×10^-1^ D) Representative image of ER-released SNB-1::SBP::eGFP (green) and SNG-1::mScarlet (magenta) containing dynamic compartments after 20 min post-biotin feeding in the PLM cell body. The yellow arrow indicates ER-released SNB-1::SBP::eGFP containing dynamic compartments containing SNG-1::mScarlet. Scale bar 5 µm (large panel with ROI), 1 µm (time series panels). E) Violin plot showing the fraction of ER-released SNB-1::SBP::eGFP containing vesicles co-migrating with SNG-1::mScarlet in the cell body of PLM neurons at 20 min post-biotin feeding. Number of animals = 15, Number of SNB-1::SBP::eGFP compartments = 103.

*C. elegans* expressing SNB-1::SBP::eGFP grown in biotin-free conditions have SNB-1::SBP::eGFP trapped in the ER. This is seen by the overlap between SNB-1::SBP::eGFP and mCherry::SP-12 marked ER (S2A). Some of the SNB-1::SBP::eGFP punctate are present adjacent with the medial Golgi marker Man-II::mCherry 10-15 min post-biotin feeding (S2B). Consistent with previous studies (11, 38, 39), SNB-1::SBP::eGFP might transit swiftly through the Golgi, suggesting that SNB-1::SBP::eGFP traffic through the Golgi in a manner consistent with the conventional ER–Golgi route. ER-released SNB-1::SBP::eGFP containing vesicles present in the proximal major neurite and minor neurite 20 min post biotin feeding (S1E and S1F). These experiments suggest that SNB-1::SBP::eGFP is restricted in the ER in the biotin auxotroph condition, and released from the ER after biotin addition and follows the conventional ER–Golgi trafficking route.

We investigated whether ER-released SNB-1::SBP::eGFP traffics through the post-Golgi SV-lysosomal intermediate compartments in the cell body. We performed dual-colour live imaging of SNB-1::SBP::eGFP with the lysosomal protein CTNS-1 (cystinosin lysosomal cystine transporter) in the cell body at various time points following biotin feeding. We quantified the fraction of moving SNB-1::SBP::eGFP co-migrating with CTNS-1::mCherry compartments in the cell body post-biotin feeding. After feeding biotin for 20 min, SNB-1::SBP::eGFP dynamic vesicles are present, but none of them contains CTNS-1::mCherry in the cell body. After 45 minutes of biotin feeding, a fraction of ER-released SNB-1–containing vesicles co-migrates with CTNS-1::mCherry (Fig 1B and 1C). Consistently, ER-released SNB-1::SBP::eGFP vesicles did not overlap with the lysosomal membrane protein LMP-1 at 20 min post-biotin feeding. However, 45 min post-biotin feeding, ER-released SNB-1::SBP::eGFP vesicles show clear co-migration with LMP-1::mScarlet (S2C and S2D).

Since pre-SVs that exit the cell body contain multiple SVPs together, we assess if ER released SNB-1::SBP::eGFP containing vesicles after 20 min post biotin feeding also have other synaptic vesicle proteins in the cell body. We performed dual-colour live imaging in the PLM neuron’s cell body and found that after 20 min of biotin feeding, a fraction of ER-released SNB-1::SBP::eGFP vesicles co-migrate with SNG-1::mScarlet (Fig 1D and 1E). ER-released SNB-1 containing moving vesicles appear in the cell body 20 minutes post biotin feeding, but are only seen with lysosomal protein at 45 minutes post biotin feeding. Overlap with ER-released SNB-1 with SNG-1 20 min post biotin feeding suggests that ER-released SNB-1::SBP::eGFP containing vesicles can also contain other synaptic vesicle proteins or traffic through compartments enriched with these proteins.

Since SV proteins have been reported to transit via the plasma membrane in the neurons (11, 12), we next tested whether ER-released SNB-1 transits through the plasma membrane before fusing with lysosomal protein-containing compartments. To block endocytosis from the plasma membrane, we used an *unc-11*/AP180 mutant (40). In *unc-11(e47)* mutant animals, the anterograde and retrograde flux of ER-released SNB-1 present in the axon is similar to wild-type animals (S3A and S3B). ER-released SNB-1 vesicles also show plasma membrane localisation in the cell body of PLM neuron 45 min post-biotin feeding in *unc-11(e47)* mutant animals (S3C), resembling the plasma membrane localisation pattern observed for plasma membrane-targeting proteins (41, 42). In *unc-11(e47)* mutant animals, ER-released SNB-1 does not overlap with CTNS-1 at 20 min post-biotin feeding, but overlaps with CTNS-1 at 45 min post-biotin feeding, similar to wild-type animals (Fig 1C, S3D). This suggests that the overlap of ER-released SNB-1 with CTNS-1 is not dependent on clathrin-mediated endocytosis from the plasma membrane. Together, these data suggest that while a fraction of ER-released SNB-1 can fuse with the plasma membrane and be endocytosed, a fraction reaches CTNS-1-positive compartments through a direct intracellular route, independent of plasma membrane transit.

### ER-released SNB-1 containing pre-SVs sequentially overlaps with SV and lysosomal proteins in the axon

Biogenesis of pre-SVs is thought to occur in the soma of neurons, where they enter the axon and are transported along the neuronal process. Unlike the cell body, where trafficking occurs across multiple planes, the axon provides a linear compartment where anterogradely moving vesicles are destined for synapses, making it better suited to resolve SVP trafficking routes and kinetics. ER-released SNB-1-containing vesicles are absent in the proximal process of PLM neuron when grown on biotin auxotroph bacteria without being fed biotin, but appear along the axon earliest at 15 minutes and appreciably at 20 minutes post-biotin feeding (S2E). Approximately 82% of ER-released SNB-1-containing vesicles move anterogradely in the proximal neuronal process (S2F), consistent with the previous observations using SNB-1::RUSH::GFP in cultured hippocampal neurons (11).

To examine whether the pre-SVs containing ER released SNB-1::SBP::eGFP contained lysosomal proteins in the axon, we performed dual-colour live imaging of SNB-1::SBP::eGFP with CTNS-1::mCherry in the proximal neuronal process. ER-released SNB-1::SBP::eGFP vesicles, coming out into the neuronal process, at 20 min post-biotin feeding, do not contain CTNS-1::mCherry, but after 45 min post-biotin feeding, the ER-released SNB-1::SBP::eGFP vesicles also contain CTNS-1::mCherry (Fig 2A and 2B). The kinetics of ER-released SNB-1::SBP overlap with CTNS-1 are similar in the cell body and the axon. SNB-1::eGFP co-migrates with CTNS-1 in the major neurite of PLM neurons, consistent with a previous study (17).

**Figure 2:**
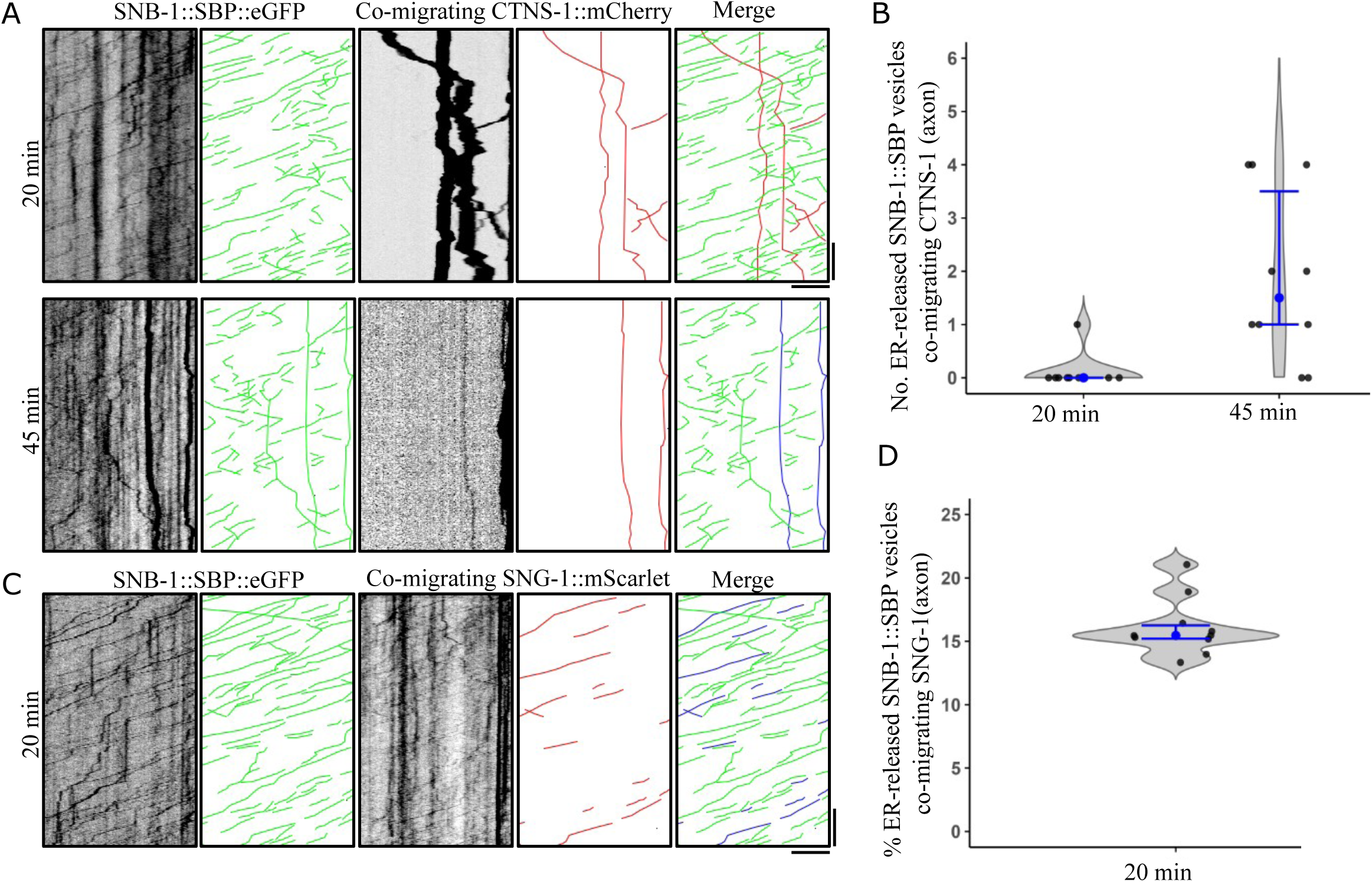
ER-released SNB-1 traffics through lysosomal-dependent and independent routes in PLM neurons. A) Dual colour imaging and kymographs for ER-released SNB-1::SBP::eGFP and CTNS-1::mCherry in the proximal major neurite of PLM neuron after 20 min and 45 min post-biotin feeding. (Green line represents SNB-1::SBP::eGFP, Red line indicates CTNS-1::mCherry, blue line indicates co-migration of SNB-1::SBP::eGFP and CTNS-1::mCherry). Scale bar x-axis 5 µm, y-axis 10 sec. B) Violin plot showing the number of ER-released SNB-1::SBP::eGFP containing vesicles co-migrating with CTNS-1::mCherry in the proximal major neurite of PLM neuron at 20 min and 45 min post-biotin feeding. Number of animals = 10, Number of SNB-1::SBP::eGFP compartments ≥ 1761. C) Dual colour imaging and kymographs for ER-released SNB-1::SBP::eGFP and SNG-1::mScarlet in the proximal major neurite of PLM neuron after 20 minutes post-biotin feeding. (Green line represents SNB-1::SBP::eGFP, red line indicates SNG-1::mScarlet, blue line indicates co-migration of SNB-1::SBP::eGFP and SNG-1::mScarlet). Scale bar x-axis 5 µm, y-axis 10 sec. D) Violin plot showing the number of ER-released SNB-1::SBP::eGFP containing vesicles co-migrating with SNG-1::mScarlet in the proximal major neurite of PLM neuron at 20 min and 45 min post-biotin feeding. Number of animals = 10, Number of SNB-1::SBP::eGFP compartments = 2050.

This suggests that ER-released SNB-1::SBP::eGFP containing pre-SVs can take two different trafficking routes to form, one route is the faster route that does not contain lysosomal proteins.

Another slower route involves the SV-lysosomal intermediate compartment, although these could also arise from the fusion of SNB-1 moving retrogradely with the CTNS-1 compartment.

Transmembrane SV proteins co-transport in the axon (16, 18), and around forty percent of total SNB-1::eGFP containing vesicles co-migrate with SNG-1::mScarlet (S5D).

We also assessed whether ER-released SNB-1::SBP::eGFP, containing pre-SVs, also contains other synaptic vesicle proteins in the neuronal process. After 20 min post-biotin feeding, approximately 17 percent of ER released SNB-1::SBP::eGFP contains another SV protein, SNG-1, in the proximal major neurite of PLM neurons (Fig 2C and 2D).

Together, these data indicate that ER-released SNB-1-containing vesicles take at least two distinct trafficking routes. Vesicles present in the proximal neuronal process at 20 minutes co-migrate with the SV protein SNG-1 but are devoid of the lysosomal marker CTNS-1, reflecting a fast, lysosome-independent route. A second population emerges at 45 minutes and acquires CTNS-1, reflecting a slower route through SV-lysosomal intermediate compartments. These observations are consistent with the existence of at least two kinetically distinct pre-SV populations, though the precise point of divergence in the secretory pathway remains to be determined.

### ER-released SNB-1 vesicles show differential recruitment of RAB-27 and RAB-3

Synaptic vesicles (SVs) contain various RABs, which are peripherally associated membrane proteins (2, 5, 6). Two specific RAB proteins, RAB-3 and RAB-27, play a role in the exocytosis of SVs at the synapse (30). However, it remains unknown at what stage during pre-SV biogenesis these RABs are acquired, and whether newly synthesised SVPs already associate with RAB-3 and RAB-27 as they exit the ER and traffic through the axon.

To test whether RAB-27 is present on pre-SVs, we examined the co-migration of RAB-27 with the SV protein SNB-1 in axons and found that 8% of SNB-1::GFP co-migrates with mCherry::RAB-27 (Fig 3A and 3B). Notably, ER-released SNB-1 containing vesicles present in the axon 20 minutes post-biotin feeding showed a similar percentage of RAB-27 co-migration in the axon (8%) (Fig 3A and 3B).

**Figure 3:**
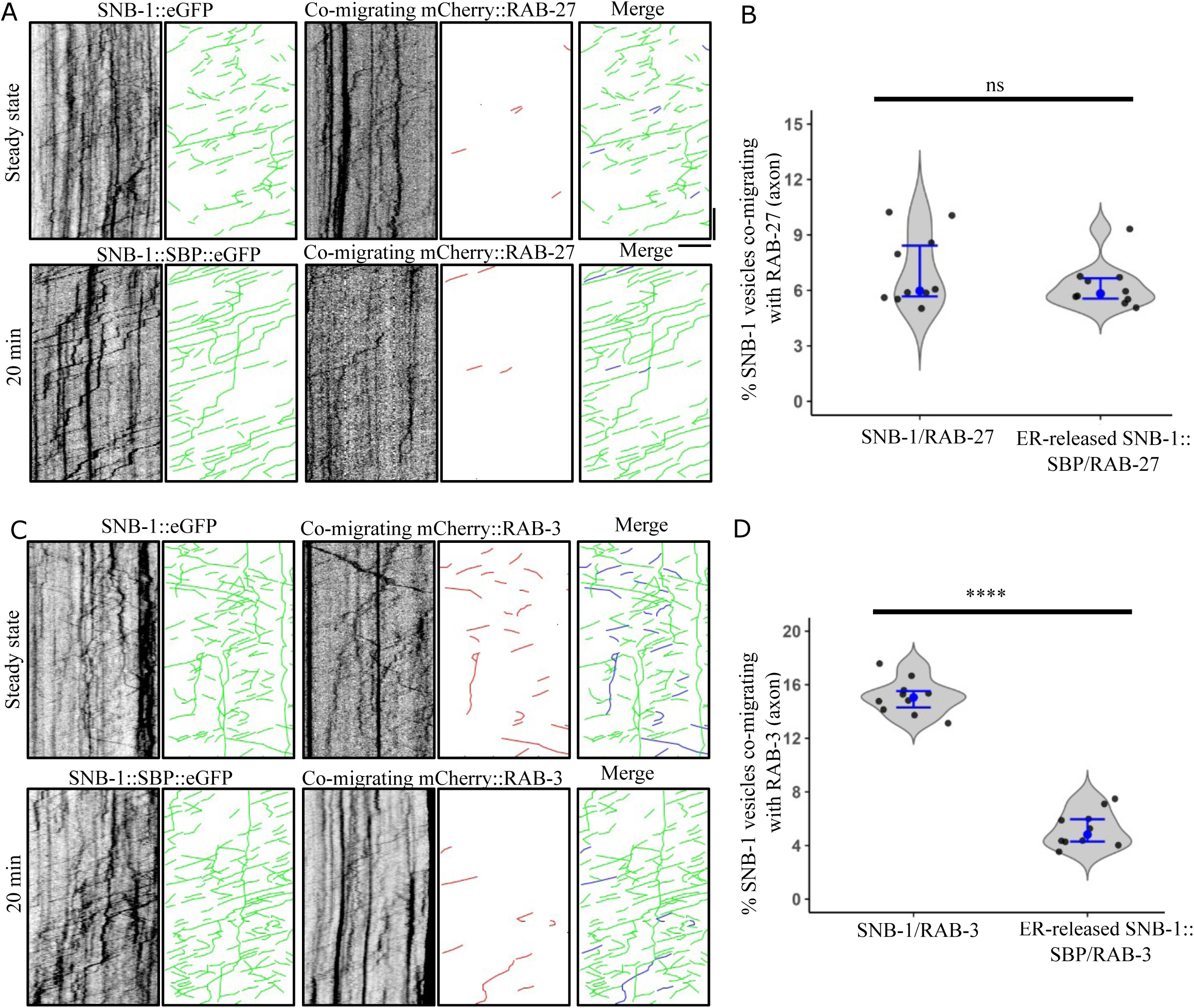
RAB-3 and RAB-27 show differential recruitment to ER-released SNB-1 vesicles. A) Dual colour imaging and kymographs of SNB-1::eGFP and mCherry::RAB-27 in steady state, and ER-released SNB-1::SBP::eGFP and mCherry::RAB-27 in the proximal major neurite of PLM neuron after 20 minutes post-biotin feeding. Scale bar x-axis 5 µm, y-axis 10 sec. B) Violin plot showing percentage of SNB-1::SBP and ER-released SNB-1::SBP::eGFP containing vesicles co-migrating with mCherry::RAB-27 in the proximal major neurite of PLM neuron. (Green line represents SNB-1 containing vesicle, red line indicates mCherry::RAB-27, blue line indicates co-migration of SNB-1 and mCherry::RAB-27) Number of animals = 10, Number of SNB-1::eGFP = 2010, Number of ER-released SNB-1::SBP::eGFP =1584. Mann-Whitney Test, *p*-value = 4.72×10^-1^ C) Dual colour imaging and kymographs of SNB-1::eGFP and mCherry::RAB-3 during steady state, and ER-released SNB-1::SBP::eGFP and mCherry::RAB-3 in the proximal major neurite of PLM neuron after 20 min post-biotin feeding. Scale bar x-axis 5 µm, y-axis 10 sec. D) Violin plot showing percentage of SNB-1::eGFP and ER-released SNB-1::SBP::eGFP containing vesicles co-migrating with mCherry::RAB-3 in the proximal major neurite of PLM neuron. (Green line represents SNB-1 containing vesicle, red line indicates mCherry::RAB-3, blue line indicates co-migration of SNB-1 and mCherry::RAB-3) Number of animals = 10, Number of SNB-1::eGFP = 3292, Number of ER-released SNB-1::SBP::eGFP =1779. Two-sample t Test, *p*-value = 2.48×10^-12^

We next examined RAB-3 co-migration in the axon. During steady state, approximately 14-15% of SNB-1-containing vesicles co-migrated with RAB-3 in axons. Surprisingly, only 6% of ER-released SNB-1 vesicles contained mCherry::RAB-3, significantly lower than steady-state levels (Fig 3C and 3D).

Together, the data indicate that ER-released SNB-1 vesicles present in the axon can carry RAB family proteins associated with SVs, but the two RABs show distinct recruitment. RAB-27 is present on a subset of SNB-1 vesicles already present on a subset of SNB-1 vesicles at early timepoints post-ER release, whereas RAB-3 could be recruited later.

### SAM-4, a BORC subunit, regulates both the kinetics of SV–lysosomal intermediate formation and the protein composition of pre-SVs

Our data suggest that SNB-1, once released from the endoplasmic reticulum (ER), can take an endo-lysosomal dependent or independent route. However, what regulates the trafficking kinetics of these routes and the composition of the resulting pre-SVs remains unknown. SAM-4/Myrlysin, a subunit of the BORC complex, is a candidate regulator: SAM-4 regulates the position of lysosomal protein but not the synaptic vesicles in the mammalian neurons (18). In *C. elegans* neurons, *sam-4/Myrlysin* regulate the localization of lysosomal proteins and synaptic vesicle protein, restricts SV-lysosomal compartments to the cell body, and reduces SV protein, RAB-3 flux into the axon (21, 25, 27). These observations suggested that *sam-4* may act on the SV-endolysosomal trafficking pathway.

We examined the co-migration of ER-released SNB-1::SBP::eGFP with CTNS-1::mCherry at various time points following biotin feeding in *sam-4*(*0*) mutant animals. In wild-type animals, ER-released SNB-1 vesicles lack CTNS-1 at this early time point (20 minutes) (Fig 1C and 4A). In contrast, in *sam-4* mutant animals, ER-released SNB-1 containing vesicles also contain CTNS-1 at 20 minutes post-biotin feeding (Fig 4A and 4B). This suggests that loss of SAM-4 shifts the kinetics of CTNS-1 co-migration with ER-released SNB-1 vesicles, with SV-lysosomal intermediate compartments forming earlier than in wild type.

**Figure 4:**
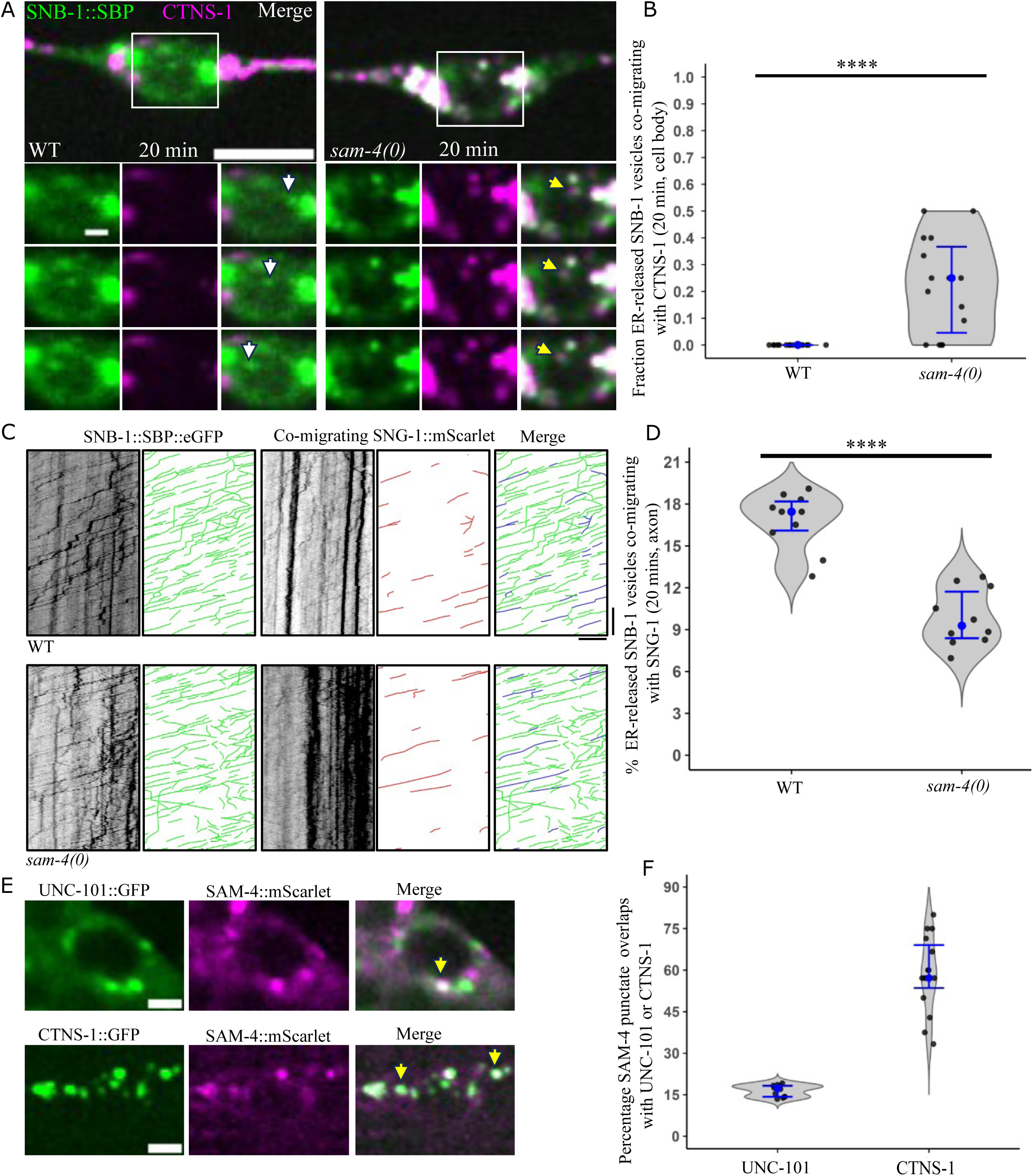
SAM-4 regulates the SV-lysosome dependent and independent routes of pre-SV biogenesis. A) Representative image of ER-released SNB-1::SBP::eGFP (green) and CTNS-1::mCherry (magenta) containing dynamic compartments after 20 min post-biotin feeding in the PLM cell body in wild type and *sam-4*(*0*). The white arrow indicates ER-released SNB-1::SBP::eGFP containing dynamic compartments lacking CTNS-1. The yellow arrow indicates ER-released SNB-1::SBP::eGFP containing dynamic compartments containing CTNS-1::mCherry. Scale bar 5 µm (large panel with ROI), 1 µm (time series panels). B) Violin plot showing fraction of ER-released SNB-1::SBP::eGFP containing vesicles co-migrating with CTNS-1::mCherry in the cell body of PLM neuron in wild type and *sam-4*(*0*) at 20 min post biotin feeding. Number of animals =15, Number of SNB-1::SBP::eGFP ≥ 71. Statistical test: Mann-Whitney Test, *p*-value = 7.38×10^-5^ C) Dual colour imaging and kymographs for ER-released SNB-1::SBP::eGFP and SNG-1::mCherry in the proximal major neurite of PLM neuron after 20 min and 45 min post-biotin feeding. Scale bar x-axis 5 µm, y-axis 10 sec. D) Violin plot showing the number of ER-released SNB-1::SBP::eGFP containing vesicles co-migrating with SNG-1::mScarlet in the proximal major neurite of PLM neuron at 20 min post-biotin feeding. Number of animals =10, Number of SNB-1::SBP::eGFP ≥ 1627 Statistical test: Two sample t Test, *p*-value= 4.98×10^-7^ E) Representative imaging showing the overlap of SAM-4::mScarlet (magenta) with UNC-101::GFP (green) and CTNS-1::GFP (green) in wild-type animals. The yellow arrow indicates the overlap between SAM-4::mScarlet with UNC-101::GFP or CTNS-1::mCherry. Scale bar 2 μm. No. of animals ≥ 12, No. of SAM-4 punctate ≥ 142.

Since the trafficking kinetics of SNB-1 co-migrating with CTNS-1 is altered, we hypothesized that SAM-4 might act at the Golgi to regulate the trafficking of SVPs. To test this, we examined the localisation of SAM-4::mScarlet with UNC-101::GFP, the μ subunit of the AP-1 complex that localises to the Golgi (36, 43). We observed that 18% of SAM-4::mScarlet puncta overlapped with UNC-101::GFP, suggesting that a subset of SAM-4 puncta may associate with the Golgi (Fig 4E and 4F). Since we know, in *sam-4* mutants, the overlap of SV-lysosomal proteins in the cell body increases (21), we checked the localisation of SAM-4 with CTNS-1 compartments. We found that SAM-4 also co-localises with CTNS-1 compartments (Fig 4E and 4F). Interestingly, while SAM-4 did not co-localise with the AP-3 complex subunit APB-3::GFP, approximately 15% of SAM-4 puncta were juxtaposed to AP-3-positive compartments (S5A and S5B), suggesting spatial proximity between SAM-4 and AP-3.

Since a subset of SAM-4 puncta associates with the Golgi, we tested whether it regulates pre-SV cargo composition entering the axon post 20 min biotin feeding. Under steady-state conditions, *sam-4* mutants showed significantly increased co-migration of SNB-1::GFP and SNG-1::mScarlet compared to wild-type animals in the proximal neuronal process (S5D) (WT: 40% vs *sam-4*: 50 %). By contrast, when we examined ER-released SNB-1::SBP::eGFP vesicles entering the proximal neuronal process at 20 minutes post-biotin feeding, we observed that the percentage of ER-released SNB-1 vesicles containing SNG-1::mScarlet was significantly reduced in *sam-4* mutants compared to wild-type animals (Fig 4C and 4D)(WT: 17% vs *sam-4*: 7 %). The steady-state increase in SNB-1 and SNG-1 co-migration in *sam-4* mutants likely reflects accumulated changes in cargo sorting over time, rather than increased co-packaging during biogenesis. The RUSH data suggest that SAM-4 is required for the proper co-packaging of SNG-1 with SNB-1 into nascent pre-SVs at the early biosynthetic route. Importantly, the total flux of ER-released SNB-1::SBP::eGFP vesicles into the proximal process remained unchanged between wild-type and *sam-4* mutants at this timepoint (S5C), indicating that SAM-4 regulates the protein composition of ER-released SNB-1 vesicles without altering the overall rate of vesicle entry into the axon via early trafficking route.

## Discussion

Accurate delivery of synaptic vesicle transmembrane proteins to the synapse requires their incorporation into pre-SVs in the cell body and transport along the neuronal process. Using the RUSH system in *C. elegans* PLM mechanosensory neurons, we showed that ER-released SNB-1 follows at least two kinetically distinct routes to the axon: a lysosome-independent pathway and a pathway dependent on SV–endolysosomal intermediate compartment. Although a fraction of ER-released SNB-1 can reach the plasma membrane and undergo endocytosis, consistent with the previous studies (11, 12), the colocalization of ER-released SNB-1 with lysosomal proteins occurs independent of plasma membrane transit. We further showed that the BORC subunit SAM-4/Myrlysin regulates both the kinetics of SV–lysosomal intermediate formation and the protein composition of pre-SVs entering the axon. Together, these findings suggest the heterogeneity underlying pre-SV biogenesis and provide a framework for understanding how distinct trafficking pathways and their regulators contribute to the fidelity of presynaptic protein delivery.

Studies across multiple model systems have established that SVPs and lysosomal proteins are found together in the same transport carriers, both in the neuronal soma and along axons (14–18, 21). A recent study using the RUSH system in hippocampal neurons suggests that ER-released lysosomal protein can co-migrate with SV proteins in axons, post one hour after biotin addition, suggesting that SV–lysosomal compartments can arise from the biosynthetic route (19). Our work using the RUSH system in *C. elegans* PLM neurons supports a model in which ER-released SNB-1 is sorted into two kinetically distinct pre-SV biogenesis routes: a lysosomal independent route, in which pre-SVs travel to the axon without transiting a lysosomal intermediate, and another route, in which SNB-1 passes through an SV–endolysosomal intermediate compartment. We propose that the lysosome-independent route allows neurons to rapidly replenish the presynaptic terminal with SV proteins during periods of high synaptic activity. The lysosome-involving route, in contrast, may serve a quality-control function, where the SV–lysosomal intermediate compartment acts as an additional sorting station. However, the precise origin of the SV–endolysosomal intermediate compartment itself remains unclear; it could arise either through the formation of vesicle at the Golgi that simultaneously incorporates SV and lysosomal proteins, or through the fusion of SNB-1-containing vesicles with a pre-existing lysosomal protein-containing compartment. Distinguishing between these possibilities will be an important goal for future work.

Previous study in hippocampal neurons suggests that synaptic vesicle (SV) proteins may utilise the plasma membrane (PM) as a reservoir; however, this conclusion is based on accumulation measured over a six-day period, which limits temporal resolution. In *unc-11* mutants, flux of ER-released SNB-1 vesicles is similar to WT, suggesting that vesicles emerging in the early route in the axon come directly without plasma membrane transit in the *C. elegans* neuron.

To determine whether SV–lysosomal overlap could arise through plasma membrane transit and endocytosis, we examined ER-released SNB-1 trafficking in *unc-11/AP180* mutants, which are deficient in clathrin-mediated endocytosis (40). The kinetics of CTNS-1 overlap and the flux of ER-released SNB-1 vesicles were both similar to wild type (Fig 1C and S3D), indicating that pre-SVs reach the axon and enter lysosomal compartments through a direct intracellular route. Although a fraction of ER-released SNB-1 was observed at the plasma membrane in *unc-11* mutants (S3C), consistent with studies in neuroendocrine cells and hippocampal neurons (8, 10–12), this route does not represent the predominant pathway for SV-lysosomal intermediate formation in *C. elegans* neurons. Previous work in hippocampal neurons has suggested that SV proteins may utilise the plasma membrane as a reservoir; however, this conclusion is based on protein accumulation measured over a six-day period, which limits temporal resolution (11), the unaltered flux of ER-released SNB-1 vesicles in *unc-11* mutants in our system indicates that vesicles entering the axon in the early route do so directly, without plasma membrane transit, in *C. elegans* neurons.

The differential recruitment of RAB GTPases to ER-released SNB-1 vesicles provides further insight into pre-SV biogenesis. The early presence of RAB-27 and the delayed recruitment of RAB-3 on newly synthesised SNB-1 mirror observations in other secretory systems; for example, during sperm acrosomal exocytosis, Rab27 acts upstream and is activated first, subsequently promoting Rab3 activation (44). Together, our findings suggest that RAB-3 and RAB-27, despite both being associated with mature SVs, are recruited to pre-SVs through temporally distinct manner.

A central question raised by our two-route model is what molecular machinery governs the decision between the direct and indirect routes. Our data suggest that SAM-4 plays a role in regulating SV–lysosomal compartment formation and pre-SV cargo composition. In sam-4 loss-of-function mutants, ER-released SNB-1 vesicles already contain CTNS-1 at 20 minutes post-biotin feeding, suggesting that loss of SAM-4 accelerates SV–lysosomal overlap, shifting what is normally a slow route into an earlier event. *sam-4* mutants show altered protein composition of ER-released SNB-1 vesicles entering the axon (Fig 4C and 4D), without any change in the total flux of ER-released SNB-1 vesicles into the axon (S5C). This indicates that SAM-4 regulates the inclusion of SV proteins into pre-SV carriers, rather than simply controlling the rate of vesicle export from the soma.

To understand where in the secretory pathway SAM-4 acts, we examined its localisation. The partial localisation of SAM-4 to the Golgi, marked by UNC-101/AP-1, suggests that SAM-4 may act at or near the TGN to regulate the early sorting of SVPs. This is further supported by the spatial proximity of SAM-4 puncta to APB-3/AP-3-positive compartments, which are known to regulate SV-lysosomal protein segregation. Although SAM-4 did not directly co-localize with AP-3, its juxtaposition suggests a functional relationship at the Golgi-endosomal interface, where the decision between lysosomal and direct axonal routes may be made.

We note that the RUSH system synchronises an artificially retained protein pool, and the observed trafficking kinetics may not perfectly recapitulate endogenous SNB-1 trafficking.

Despite this caveat, our data suggest that the heterogeneity of pre-SVs observed under steady-state conditions is not simply stochastic, but reflects the existence of distinct, regulated biogenesis routes. An important future direction will be to determine whether these two routes deliver functionally distinct SV populations to the synapse, and whether the balance between them is modulated by neuronal activity or developmental state.

## Acknowledgments

We thank Dr. Souvik Modi for making the construct *mec-4P*::mCherry::SP12 and the transgenic line *tbEx323*. We thank Dr. Amal Mathew for making the construct *mec-4P*::mCherry::RAB-27 and for providing comments on the manuscript. We thank Dr. Michael L. Nonet for providing strain *jsSi2013 tbSi521*. Some strains were provided by the CGC, which is funded by NIH Office of Research Infrastructure Programs (P40 OD010440). Research in Sandhya Koushika’s laboratory is supported by grants from the Department of Atomic Energy, Government of India (DAE; OM no. 1303/2/2019/R&D-II/DAE/2079; Project identification number RTI4003 dated 11.02.2020). During the preparation of this manuscript, the authors used Grammarly for grammar and spell-checking, Inkscape for figure preparation, and Claude Sonnet 4.5 (Anthropic) for refining analysis code.

## Methods

### Maintenance and growth of *C. elegans* strains

All *Caenorhabditis elegans* strains were maintained at 20 °C on standard Nematode Growth Medium (NGM) agar plates seeded with Escherichia coli OP50 (45). Before seeding, *E. coli* OP50 was cultured overnight at 37 °C in LB broth.

For biotin-free conditions, the biotin auxotroph *E. coli* MG1655*bioB:kan* was grown in M9 minimal medium using glucose as a carbon source and seeded onto NGM agar plates that do not contain peptone. Adult worms grown on OP50 *E. coli* were transferred to plates containing biotin auxotroph bacteria. F1 progeny raised on biotin auxotroph bacteria were used for imaging. Worms grown in biotin-free conditions were fed with 50 mM D-biotin for different times before imaging.

### Generation of transgenic lines

Transgenic lines of *Caenorhabditis elegans* were generated by following the standard microinjection technique (46) by using an Olympus IX53 inverted microscope (4X, 20X and 40X air objectives), a Narishige M-152 micromanipulator, and an Eppendorf FemtoJet 2 microinjector. The progenies that inherited and stably expressed the extrachromosomal transgene with less than 50% transmission were UV-irradiated to integrate the DNA and generate transgenic lines. These lines were backcrossed at least six times before use. The strains used in the study are listed in the supplementary table S1.

### Image acquisition

Larval stage 4 animals were anesthetized using 5mM tetramisole and mounted on 5% agar pads. Live imaging in the cell body and the axon was performed using a 100X/1.40 NA oil objective lens with 2 x 2 binning using Olympus IX83 fitted with a Yokogawa CSU-W1 SoRa scan unit, excited with 488 nm and/or 561 nm laser imaged using a Prime BSI sCMOS camera. The effective pixel size at the specimen plane was 0.13 µm. Dual-colour movies were acquired sequentially at 120 ms exposure time for approximately 3 minutes.

Z-stack images corresponding to panels S2A and S2B were acquired on a Zeiss LSM 880 confocal microscope (Carl Zeiss, Germany) equipped with a Plan-Apochromat 63×/1.40 NA oil immersion objective. Two channels were imaged sequentially: EGFP (excitation 488 nm) and mCherry (excitation 561 nm), using PMT detectors. Z-stacks were collected with a step size of 0.30 µm.

### Image Analysis

Fiji-ImageJ was used for image processing, analysis, and figure preparation (47).

### Co-migration in the cell body

Both images were opened in ImageJ, and vesicle movement was tracked frame by frame using the “synchronize window” tool in ImageJ.

Co-migration of CTNS-1::mCherry or SNG-1::mScarlet with dynamic ER-released SNB-1::SBP::eGFP was defined as instances where both proteins overlapped and moved along similar trajectories in the cell body. A vesicle was counted as moving if it displaced a minimum of 5 pixels. Co-migration of CTNS-1::mCherry or SNG-1::mScarlet with dynamic ER-released SNB-1::SBP::eGFP was defined as instances where both proteins overlapped and moved together along the same trajectory for its entire duration.

### Kymograph analysis in the axon

To calculate the co-migration of two proteins in the axon, kymographs were generated from a segmented line drawn over the same region of the movie in both channels using the ImageJ plugin “Multi Kymograph”. Both kymographs were opened, and visually distinct trajectories were manually marked using the segmented lines tool in ImageJ. A co-migration event was defined as an instance where the sloped lines from both channels overlapped.

Fraction of ER-released SNB-1::SBP::eGFP co-migrating with another protein = Co-migration events/ Total number of ER-released SNB-1::SBP::eGFP containing vesicles in the axon.

To determine the direction of movement of vesicle trajectories, all moving vesicles were tracked and categorised based on their direction of motion as either anterograde or retrograde. Vesicles moving away from the cell body were classified as anterograde, whereas those moving toward the cell body were classified as retrograde.

Vesicle flux was calculated as the total number of moving vesicles passing per micron per minute.

Transport parameters, including vesicle movement direction and flux, were quantified using a custom-written ImageJ macro (available at https://github.com/badal703/ImageJ-Macros).

### Rescue of snb-1(md247)

L4-stage worms were picked and placed on 35 mm plates seeded with 10 µL of OP50 *E. coli* culture and incubated for 6 hours. After incubation, freely moving worms were imaged at a rate of 1 frame per second. Worm velocity was calculated by manually tracing the paths followed by the worms for one minute.

### Quantifying the SAM-4 localisation with respect to UNC-101, APB-3 and CTNS-1

Imaging was performed on an Olympus IX83 microscope equipped with a Yokogawa CSU-W1 spinning disk unit. Z-stack images were captured using the SoRA module in cellSens software, with a 100× oil immersion objective (1.4 NA, DIC) and a Prime BSI sCMOS camera. Fluorescence excitation was achieved using 473 nm and 561 nm solid-state lasers. Images were acquired sequentially along the Z-axis with a 0.28 µm step size and a 500 ms exposure time.

To assess the spatial relationship between SAM-4::mScarlet and APB-3::GFP, a line intensity profile was drawn across puncta in both channels simultaneously using the same line. Two puncta were scored as juxtaposed if the distance between their peak intensity values was 3 pixels or less. The percentage of SAM-4 puncta juxtaposed to APB-3-positive compartments was calculated as the number of juxtaposed SAM-4 puncta divided by the total number of SAM-4 puncta analysed. The code used for peak intensity calculation is available at (https://github.com/badal703/ImageJ-Macros).

### Statistical analysis

Normality of data distributions was assessed using the Shapiro–Wilk test in Origin software. For datasets that met the assumption of normality, statistical significance was evaluated using an unpaired two-tailed t-test with unequal variances (Welch’s correction). For datasets that did not meet normality, the Mann–Whitney U test was applied.

Data are presented as violin plots with individual data points overlaid; the median is indicated by a point, and the 25th and 75th percentiles are shown as error bars. Statistical significance is denoted as follows: p < 0.05 (*), p < 0.01 (**), p < 0.001 (***), p < 0.0001 (****), and not significant (ns).

All numerical values used to generate the graphs are provided in Table S3, and all statistical comparisons are summarised in Table S4.

## Legends for supplementary documents

**Table S1:** List of strains used in this study.

**Table S2:** Provides details and procedures for the generation of the plasmids and transgenes.

**Table S3:** Original datasets used to generate the plots presented in each figure. Each sheet corresponds to an individual graph and includes the underlying raw data, organised according to figure number.

**Table S4:** Description of the statistical analyses performed, including the specific tests applied and the corresponding p-values, organised according to the figure.

