## Supplementary material for "Temporal tracking of Synaptobrevin-1 trafficking reveals SAM-4/BORC-dependent trafficking routes in *C. elegans* neurons": Movie legends

### Supplementary movie legends

**Movie 1: 3D projection of SNB-1::SBP::eGFP and MAN-II::mCherry in wild-type animals**

3D projection of ER-released SNB-1::SBP::eGFP (green) and MAN-II::mCherry (magenta) in the PLM cell body of wild-type animals 10 min post-biotin feeding. Playback speed at 2 frames per second (fps). Genotype: *tbIs457; tbEx254,* Scale: 5 µm.

**Movie 2: SNB-1::SBP::eGFP and CTNS-1::mCherry before biotin feeding in wild-type animals**

Dynamic SNB-1::SBP::eGFP (green) and CTNS-1::mCherry (magenta) in the PLM cell body of wild-type animals before biotin feeding. Imaged sequentially at 3.2 frames per second (fps), playback speed at 10 fps. Genotype: *tbIs457; tbIs381,* Scale bar: 5 µm.

**Movie 3: SNB-1::SBP::eGFP and CTNS-1::mCherry at 20 min post-biotin feeding in wild-type animals**

Dynamic SNB-1::SBP::eGFP (green) and CTNS-1::mCherry (magenta) in the PLM cell body of wild-type animals 20 min post-biotin feeding. Imaged sequentially at 3.2 frames per second (fps), playback speed at 10 fps. Genotype: *tbIs457; tbIs381,* Scale bar: 5 µm.

**Movie 4: SNB-1::SBP::eGFP and CTNS-1::mCherry at 20 min post-biotin feeding in wild-type animals**

Dynamic SNB-1::SBP::eGFP (green) and CTNS-1::mCherry (magenta) in the PLM cell body of wild-type animals 45 min post-biotin feeding. Imaged sequentially at 3.2 frames per second (fps), playback speed 10 fps. Genotype: *tbIs457; tbIs381,* Scale bar: 1 µm.

**Movie 5: SNB-1::SBP::eGFP and SNG-1::mScarlet at 20 min post-biotin feeding in wild-type animals**

Dynamic SNB-1::SBP::eGFP (green) and SNG-1::mScarlet (magenta) in the PLM cell body of wild-type animals 20 min post-biotin feeding. Imaged sequentially at 3.2 frames per second (fps), playback speed 10 fps. Genotype: *tbIs457; tbSi521,* Scale bar: 1 µm.

**Movie 6: SNB-1::SBP::eGFP before biotin feeding in wild-type animals**

Dynamic SNB-1::SBP::eGFP (green) in the PLM major neurite of wild-type animals 0 min post biotin feeding. Imaged sequentially at 3.2 frames per second (fps), playback speed 10 fps. Genotype: *tbIs457 ,* Scale bar: 5 µm.

**Movie 7: SNB-1::SBP::eGFP at 20 min post-biotin feeding in wild-type animals**

Dynamic SNB-1::SBP::eGFP (green) in the PLM major neurite of wild-type animals 20 min post-biotin feeding. Imaged sequentially at 3.2 frames per second (fps), playback speed 10 fps. Genotype: *tbIs457 ,* Scale bar: 5 µm.

**Movie 8: SNB-1::SBP::eGFP and CTNS-1::mCherry at 20 min post-biotin feeding in wild-type animals**

Dynamic SNB-1::SBP::eGFP (green) and CTNS-1::mCherry (magenta) in the PLM major neurite of wild-type animals 20 min post-biotin feeding. Imaged sequentially at 3.2 frames per second (fps), playback speed 10 fps. Genotype: *tbIs457; tbIs381,* Scale bar: 5 µm.

**Movie 9: SNB-1::SBP::eGFP and CTNS-1::mCherry at 45 min post-biotin feeding in wild-type animals**

Dynamic SNB-1::SBP::eGFP (green) and CTNS-1::mCherry (magenta) in the PLM major neurite of wild-type animals 45 min post-biotin feeding. Imaged sequentially at 3.2 frames per second (fps), playback speed 10 fps. Genotype: *tbIs457; tbIs381,* Scale bar: 5 µm.

**Movie 10: SNB-1::SBP::eGFP and SNG-1::mScarlet at 20 min post-biotin feeding in wild-type animals**

Dynamic SNB-1::SBP::eGFP (green) and SNG-1::mScarlet (magenta) in the PLM major neurite of wild-type animals 20 min post-biotin feeding. Imaged sequentially at 3.2 frames per second (fps), playback speed 10 fps. Genotype:*; tbIs457; tbSi521,* Scale: 5 µm.

**Movie 11: SNB-1::eGFP and mCherry::RAB-27 in wild-type animals**

Dynamic SNB-1::eGFP (green) and mCherry::RAB-27 (magenta) in the PLM major neurite of wild-type animals during steady state. Imaged sequentially at 3.2 frames per second (fps), playback speed 10 fps. Genotype: *tbIs457; tbEx585,* Scale bar: 5 µm.

**Movie 12: SNB-1::SBP::eGFP and mCherry::RAB-27 in wild-type animals**

Dynamic SNB-1::SBP::eGFP (green) and mCherry::RAB-27 (magenta) in the PLM major neurite of wild-type animals 20 min post-biotin feeding. Imaged sequentially at 3.2 frames per second (fps), playback speed 10 fps. Genotype: *tbIs457; tbEx585,* Scale bar: 5 µm.

**Movie 13: SNB-1::eGFP and mCherry::RAB-3 in wild-type animals**

Dynamic SNB-1::eGFP (green) and mCherry::RAB-27 (magenta) in the PLM major neurite of wild-type animals during steady state. Imaged sequentially at 3.2 frames per second (fps), playback speed 10 fps. Genotype: *tbIs457; tbIs227,* Scale bar: 5 µm.

**Movie 14: SNB-1::SBP::eGFP and mCherry::RAB-3 in wild-type animals**

Dynamic SNB-1::SBP::eGFP (green) and mCherry::RAB-27 (magenta) in the PLM major neurite of wild-type animals 20 min post-biotin feeding. Imaged sequentially at 3.2 frames per second (fps), playback speed 10 fps. Genotype: *tbIs457; tbIs227,* Scale bar: 5 µm.

**Movie 15: SNB-1::SBP::eGFP and CTNS-1::mCherry in *sam-4(0)* animals**

Dynamic SNB-1::SBP::eGFP (green) and CTNS-1::mCherry (magenta) in the PLM cell body of *sam-4(0)* animals 20 min post biotin feeding. Imaged sequentially at 3.2 frames per second (fps), playback speed 10 fps. Genotype: *tbIs457; sam-4(js415); tbIs381,* Scale bar: 1 µm.

**Movie 16: SNB-1::SBP::eGFP and SNG-1::mScarlet in *sam-4(0)* animals**

Dynamic SNB-1::SBP::eGFP (green) and SNG-1::mScarlet (magenta) in the PLM major neurite of *sam-4(0)* animals 20 min post-biotin feeding. Imaged sequentially at 3.2 frames per second (fps), playback speed 10 fps. Genotype: *tbIs457; sam-4(js415); tbSi521,* Scale bar: 5 µm.

**Movie 17: SNB-1::SBP::eGFP and CTNS-1::mCherry in *unc-11(e47)* animals**

Dynamic SNB-1::SBP::eGFP (green) and CTNS-1::mCherry (magenta) in the PLM cell body of *unc-11(e47)* animals 20 min post-biotin feeding. Imaged sequentially at 3.2 frames per second (fps), playback speed 10 fps. Genotype: *unc-11(e47) tbIs457; tbIs381,* Scale bar: 5 µm.

**Movie 18: SNB-1::SBP::eGFP and CTNS-1::mCherry in *unc-11(e47)* animals**

Dynamic SNB-1::SBP::eGFP (green) and CTNS-1::mCherry (magenta) in the PLM cell body of *unc-11(e47)* animals 45 min post-biotin feeding. Imaged sequentially at 3.2 frames per second (fps), playback speed 10 fps. Genotype: *unc-11(e47) tbIs457; tbIs381,* Scale bar: 1 µm.
