## Supplementary Figures for "Temporal tracking of Synaptobrevin-1 trafficking reveals SAM-4/BORC-dependent trafficking routes in *C. elegans* neurons"

### Supplementary Figure 1

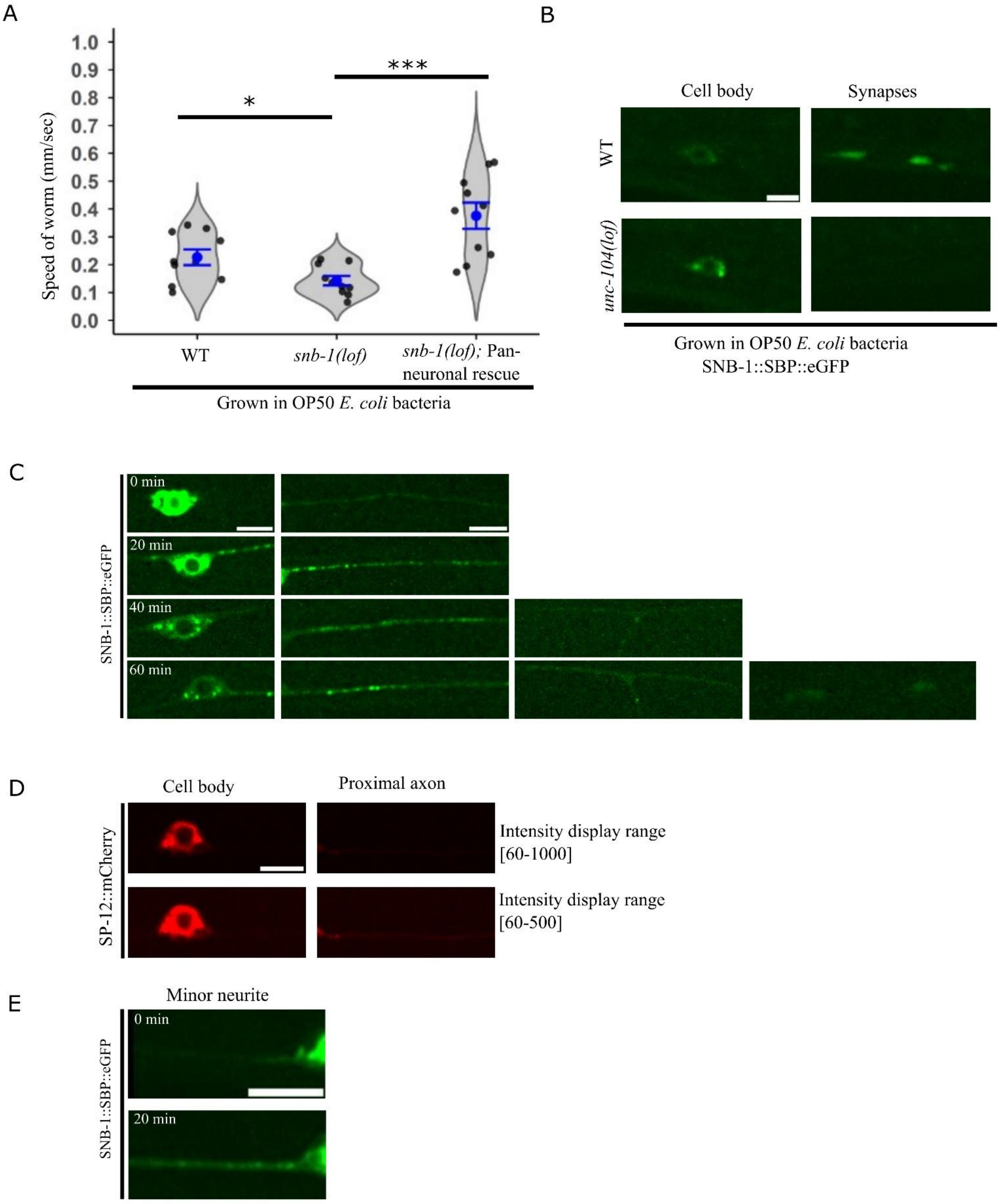

**Supplementary Figure 1**

A) Schematic representation of *C. elegans* touch receptor neurons (TRNs) (image not to scale).

B) Violin plot showing the speed of locomotion: N2, *snb-1(lof),* and *snb-1(lof)* expressing pan-neuronal SNB-1::SBP::eGFP grown on OP50. No. of animals = 10, Statistical test = Unpaired two-tailed Welch's t-test. N2 vs *snb-1(lof)* $p$-value = 0.02, *snb-1(lof)* vs *snb-1(lof)* pan neuronal overexpression $p$-value = $6.93 \times 10^{-4}$.

C) Schematic representation of the PLM touch receptor neuron regions: the minor neurite, cell body (soma), proximal axon, branch point, and synapses. Red boxes indicate the regions of interest for imaging.

D) Representative image of SNB-1::SBP::eGFP present in the cell body and at synapses in wild-type and *unc-104(e1265)* mutant animals grown on OP50 bacteria. Scale bar 5µm.

E) Localisation of SNB-1::SBP::eGFP in the PLM major neurite post-biotin feeding for different time intervals (0, 20, 40, and 60 min). Scale bar 5µm.

F) Representative image of mCherry::SP-12 present in the cell body and the proximal neuronal process in animals grown on biotin-auxotroph *E. coli*. Scale bar 5µm.

G) Representative image of SNB-1::SBP::eGFP present in the minor neurite of the PLM neuron post-biotin feeding at different time intervals (0 and 20 min). Scale bar 5µm.

### Supplementary Figure 2

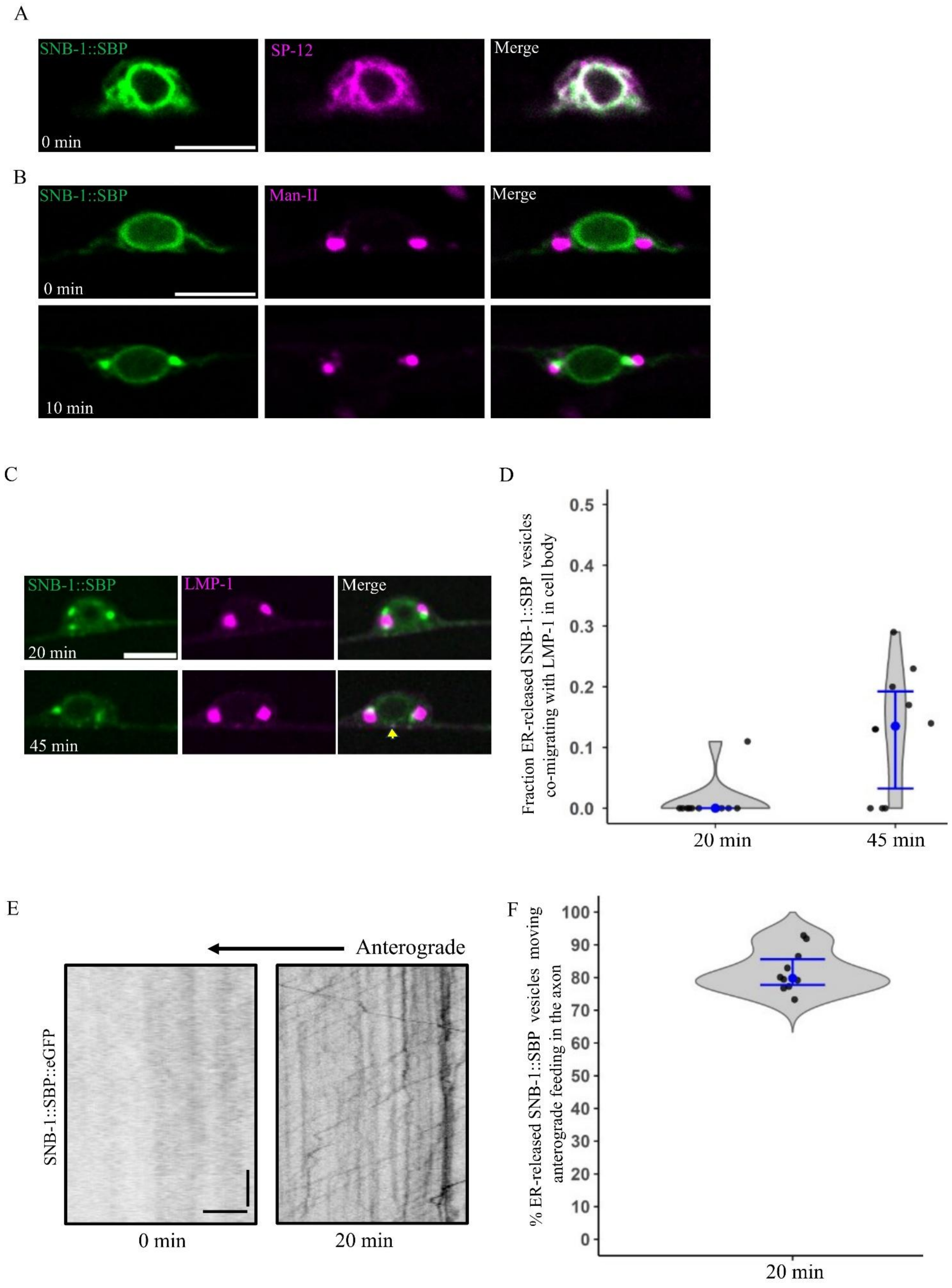

**Supplementary Figure 2**

A) Representative image of SNB-1::SBP::eGFP (green) with mCherry::SP-12 (magenta) pre-biotin feeding in the PLM cell body. Scale bar 5µm.

B) Representative image of SNB-1::SBP::eGFP (green) with Man-II::mCherry (magenta) pre-biotin feeding and 10 min post-biotin feeding in the PLM cell body. Scale bar 5µm.

C) Representative image of ER-released SNB-1::SBP::eGFP (green) and LMP-1::mScarlet (magenta) dynamic compartments after 20 min and 45 min post-biotin feeding in the PLM cell body. The yellow arrow indicates ER-released SNB-1::SBP::eGFP containing dynamic compartments containing LMP-1::mScarlet. Scale bar 5 µm.

D) Violin plot showing the fraction of ER-released SNB-1::SBP::eGFP containing vesicles co-migrating with LMP-1::mScarlet in the PLM neuron cell body at 20 min and 45 min post-biotin feeding. No. of animals = 10, No. of ER-released SNB-1::SBP::eGFP containing vesicles ≥ 64.

E) Kymograph of SNB-1::SBP::eGFP containing vesicles in the proximal major neurite of the PLM neurons at 0 min and 20 min of biotin feeding. Scale bar x-axis = 5 um, y-axis = 10 sec.

F) Violin plot showing the percentage of ER-released SNB-1::SBP::eGFP containing vesicles moving anterogradely in the proximal neuronal process 20 min post-biotin feeding. Number of animals = 10, No. of ER-released SNB-1::eGFP containing vesicles = 2457.

### Supplementary Figure 3

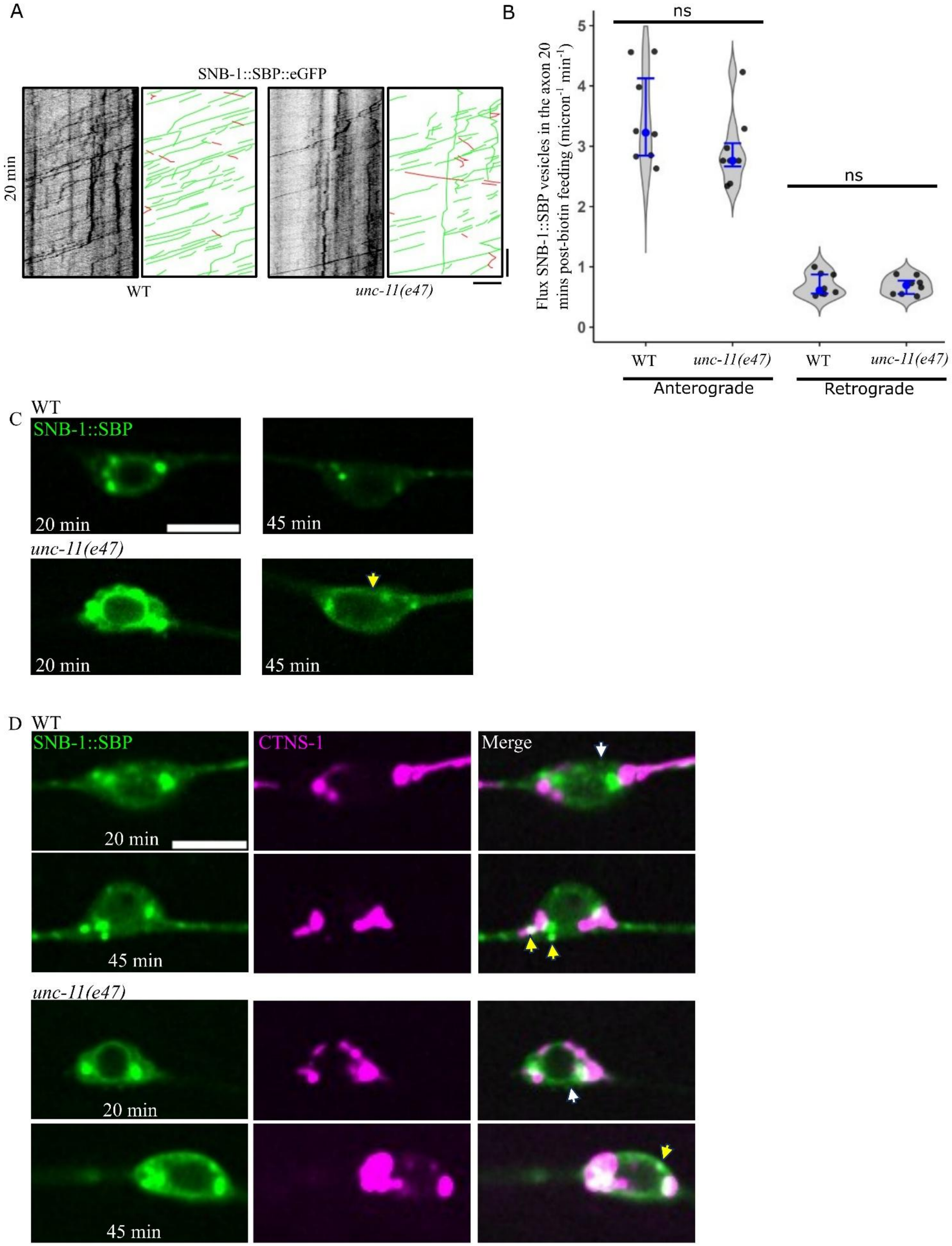

**Supplementary Figure 3**

A) Kymograph of SNB-1::SBP::eGFP containing vesicles in the proximal major neurite of the PLM neurons at 20 min post-biotin feeding in WT and *unc-11(e47)* mutant animals. Scale bar x-axis = 5 um, y-axis = 10 sec.

B) Violin plot showing the flux of ER-released SNB-1::SBP::eGFP containing vesicles moving anterogradely and retrogradely in the major neurite of the PLM neuron at 20 min post-biotin feeding in WT and *unc-11(e47)* mutant animals. No. of animals = 8, No. of ER-released SNB-1::SBP::eGFP containing vesicles ≥ 1194. Statistical test = Unpaired two-tailed Welch's t-test,
anterograde flux (WT vs *unc-11*) *p*-value = $1.4 \times 10^{-1}$, retrograde flux (WT vs *unc-11*) *p*-value = $8.6 \times 10^{-1}$.

C) Representative image of SNB-1::SBP::eGFP in the PLM cell body, 20 min and 45 min post-biotin feeding in WT and *unc-11(e47)* mutant animals. Scale bar 5µm.

D) Representative image of ER-released SNB-1::SBP::eGFP (green) and CTNS-1::mCherry (magenta) in the PLM cell body of wild-type and *unc-11(e47)* mutant animals at 20 min and 45 min post-biotin feeding. The white arrow indicates ER-released SNB-1::SBP::eGFP containing dynamic compartments that do not contain CTNS-1::mCherry. The yellow arrow indicates ER-released SNB-1::SBP::eGFP containing dynamic compartments containing CTNS-1::mCherry. Scale bar 5µm.

### Supplementary Figure 4

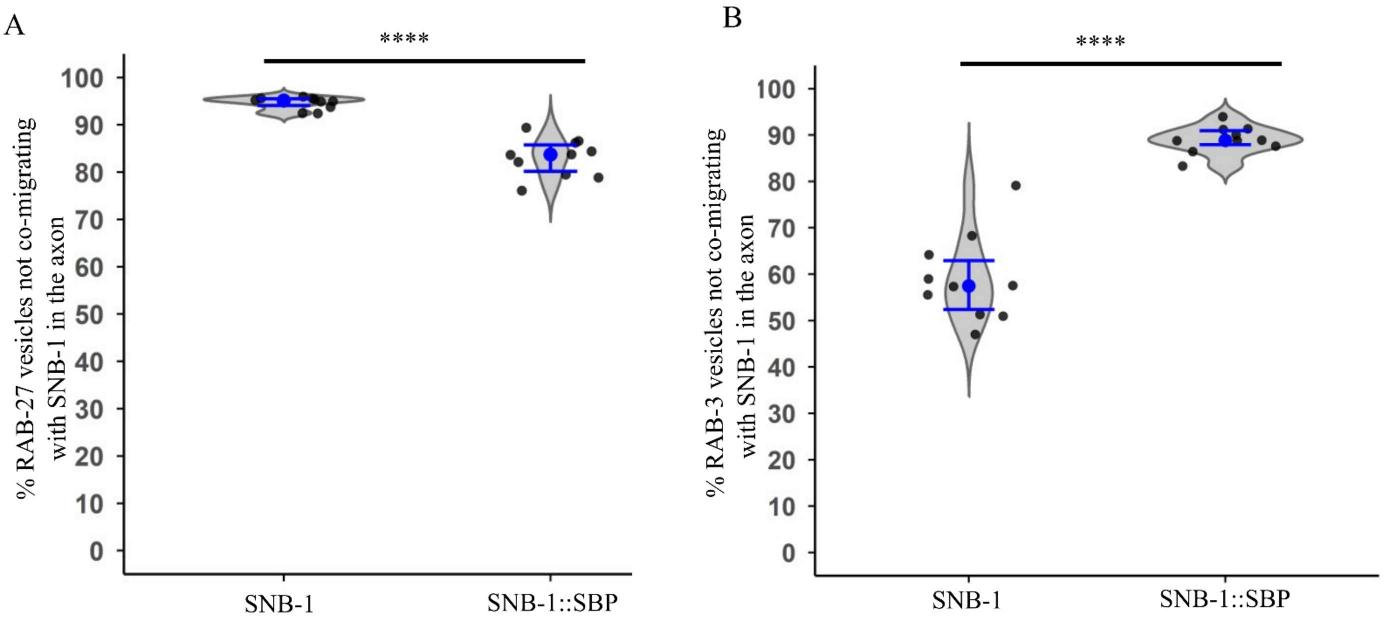


**Supplementary Figure 4**

A) Violin plot showing percentage of RAB-27 containing vesicles not co-migrating with SNB-1 or SNB-1::SBP in the proximal major neurite of the PLM neuron. Number of animals = 10, No. of mCherry::RAB-27 containing vesicles ≥ 620. Statistical test = Mann-Whitney Test. p-value = $3.29 \times 10^{-6}$

B) Violin plot showing percentage of RAB-3 containing vesicles not co-migrating with SNB-1 or SNB-1::SBP in the proximal major neurite of the PLM neuron. Number of animals = 10, No. of mCherry::RAB-3 containing vesicles ≥ 849. Statistical test = Unpaired two-tailed Welch's t-test. $p$-value = $1.37 \times 10^{-8}$.

### Supplementary Figure 5

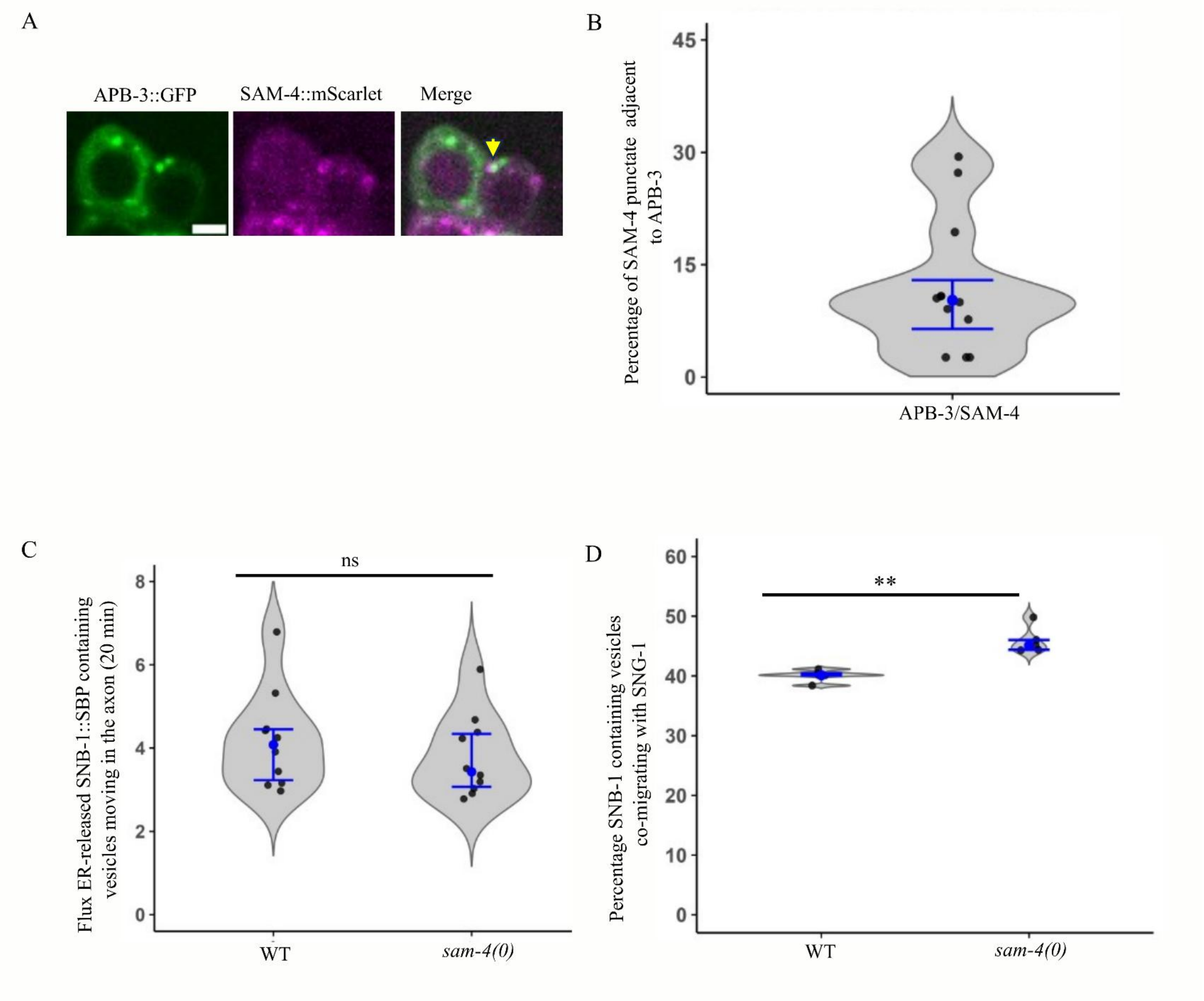

**Supplementary Figure 5**

A) Representative images showing SAM-4::mScarlet (magenta) puncta adjacent to APB-3::GFP (green) in wild-type animals. Scale bar 2 μm. The yellow arrow indicates the SAM-4::mScarlet puncta adjacent to APB-3::GFP.

B) Quantification showing percentage of SAM-4::mScarlet (magenta) puncta adjacent to APB-3::GFP (green) in wild-type animals. No. of animals = 12, No. of SAM-4 puncta = 377.

C) Violin plot showing the flux of ER-released SNB-1::SBP::eGFP containing vesicles in the wild-type and *sam-4(js415)* mutant animals 20 min post-biotin feeding. No. of animals = 10, No. of ER-released SNB-1::SBP::eGFP containing vesicles ≥ 1627, Statistical test = Unpaired two-tailed Welch's t-test *p*-value = $4.35 \times 10^{-1}$.

D) Quantification showing percentage of SNB-1::eGFP containing vesicles co-migrating with SNG-1::mScarlet in the proximal major neurite of the PLM neurons during steady state in WT and *sam-4(js415*) mutant animals. No. of animals = 5, No. of SNB-1::eGFP containing vesicles ≥ 1252, Statistical test = Unpaired two-tailed Welch's t-test. *p*-value = $2.50 \times 10^{-3}$.
