## Supplementary material for "Temporal tracking of Synaptobrevin-1 trafficking reveals SAM-4/BORC-dependent trafficking routes in *C. elegans* neurons": Table S1

| S. No. | Strain No. | Genotype | Reference |
|---|---|---|---|
| 1 | N2 | Bristol wild type | [1] |
| 2 | | *sam-4(js415) II null (0)* | [2] |
| 3 | TT2903 | *tbIs381 [mec-4p::ctns-1::mCherry (20 ng/µl), myo-2p::h2b::gfp (40 ng/µl), pBluescript SK (110 ng/µl)] X* | [2] |
| 4 | TT3007 | *tbIs414 [mec-7p::snb-1::egfp (10 ng/µl), myo-2p::mCherry (10 ng/µl), pBluescript SK (180 ng/µl)]* | [2] |
| 5 | TT3608 | *tbIs518 [rab-3p::apb-3::gfp (30 ng/µl), myo-2p::mCherry (10 ng/µl), pBluescript SK (140 ng/µl)]* | [5] |
| 6 | WEH644 | *sam-4(syb4105[sam-4::mScarlet-I::ZF1]) II (CRISPR)* | [4] |
| 7 | TT1884 | *tbIs227 [mec-4p::mCherry::rab-3]* | [5] |
| 8 | TT1893 | *oqEx [unc-101p::unc-101::gfp+pRF4]* | [6] |
| 9 | TT4021 | *tbEx585 [mec-4p::mCherry::rab-27*(TTpl730)(10ng/ul); *myo-2p::gfp::h2b*(TTpl592)(50ng/ul); *pBluescript*(TTpl542)(140ng/ul)] | This study |
| 10 | TT3177 | *tbIs457* [Integrated tbEx426 [*mec-4p::snb-1::sbp::egfp* (TTpl 707) (10ng/uL), *rab-3p::streptavidin::KDEL* (TTpl 736) (30ng/uL), *pBluescript SK* (TTpl542) (160 ng/uL)]] | This study |
| 11 | CB47 | *unc-11(e47)* I. | CGC |
| 12 | TT2619 | *tbEx323* [*pmec4::mCherry::sp-12*(TT623) (10ng/ul)+*ttx-3p::rfp* (TT541) (50ng/ul)+*pBluscriptSK*(TT542) (140ng/ul). 40% penetrance] | This Study |
| 13 | TT3395 | *tbEx488-[rab-3p::snb-1::sbp::egfp (30ng/uL), myo-2p::h2b::gfp (50ng/uL) (TTpl 592),pBlueScript (120 ng/uL) (TTpl542) Penetrance ~50 %]* | This Study |
| 14 | NM467 | *snb-1(md247)* | CGC |
| 15 | TT1555 | *tbEx254 [mec-7p::man-II::mCherry]*; *jsIs821* | [5] |
| 16 | TT3482 | *jsSi2013 tbSi521* [loxP *mec-4Sp sng-1-L-mScarlet*-C1 *tbb-2* 3' FRT3] | This Study |
| 17 | TT3028 | *pwSi222 [mec-7P::lmp-*1::*mScarlet]* | [7] |
